# Reclamation is not the primary determinant of soil recovery from oil and gas development in Wyoming sagebrush systems

**DOI:** 10.1101/546804

**Authors:** Caitlin M. Rottler, Ingrid C. Burke, William K. Lauenroth

**Affiliations:** Program in Ecology, University of Wyoming, Laramie, Wyoming, United States of America; Department of Botany, University of Wyoming, Laramie, Wyoming, United States of America; Haub School of Environment and Natural Resources, University of Wyoming, Laramie, Wyoming, United States of America

## Abstract

Dryland soils store approximately 10-15% of the world’s soil organic matter (SOM) to 1 m. Threats to carbon stocks in global dryland soils include cultivation, overgrazing, urbanization, and energy development. To limit loss of carbon from these soils, it is important to understand, first, how disturbances affect SOM and second, how SOM recovers after disturbance. In this study, we address current gaps in our understanding of the effects of oil and gas development and reclamation on SOM in the sagebrush steppe of Wyoming, a cold temperate shrub-dominated dryland. Most studies have found that soil disturbance, including from the respreading of topsoil during wellpad reclamation, is damaging to SOM stores; however, research on ~80 year old unreclaimed oil and gas wellpads found no difference in SOM between wellpads and undisturbed sites. Using a chronosequence approach and paired study design, we evaluated the effects of reclamation on SOM by comparing undisturbed sites to wellpads where reclamation activities either had or had not occurred. Our results suggest that the most important factor in recovery of SOM after disturbance in this area was not the presence or absence of reclamation, but time since wellpad abandonment and spatial heterogeneity of plants. Further study on the effectiveness of different reclamation techniques is warranted if the goal of reclamation is to aid SOM recovery and prevent further C loss from these systems.

## Introduction

Soil organic matter (SOM) represents a large pool of soil organic carbon (SOC) in ecosystems and plays an important role in the global carbon (C) cycle [1,2]. Globally, between 15-16% of soil carbon to 1m is stored in dryland ecosystems [3]. Past disturbance from land degradation and desertification has resulted in loss of carbon from these systems, causing many of them to become carbon sources to the atmosphere [3–5]. Of the carbon lost, researchers estimate that up to 2/3 could be re-sequestered through a variety of management practices designed to increase inputs and decrease outputs [3], ultimately turning them from sources of C into sinks. The ability of SOM pools to act as sources or sinks of CO_2_ depending on land use and environmental conditions has multiple implications for global change [1,6–8], and is an avenue by which responsible management can help avoid further contribution to global change.

Disturbances from a variety of land uses, including cultivation, overgrazing, urbanization, and various forms of energy development, can have both short- and long-term effects on C stored in dryland soils. One major factor in C storage is aggregate structure, especially the formation and stability of macro- and microaggregates, which bind to C and C-containing compounds and help prevent or slow their breakdown [9]. These aggregates are affected by land use, soil texture, and quality and quantity of plant inputs to the soil [10,11]. As a result of long recovery times, disturbance can be especially damaging to aggregates associated with very fine soil particle sizes.

One important source of soil disturbance in western US drylands is energy development, which has received increasing attention due to its significant and growing role in western economies [12]. In particular, the growth of the oil and gas industry has led to widespread surface soil disturbance through drilling of wells [13]. Current practices require these disturbed areas be reclaimed [14]. However, the requirements and standards, especially for oil and gas drilling, are vague and open to interpretation, resulting in reclamation that is often inefficient or ineffective [15,16]. In addition, the responsibility for enforcing reclamation laws is dependent upon land ownership. Much of the western United States falls under the jurisdiction of the Bureau of Land Management, and is therefore subject to BLM regulations [14]. As with other surface soil disturbances, drilling for oil and gas results in destruction of soil aggregates and loss of SOM and associated nutrients, which can hinder ecosystem recovery and limit the ability of the soil to store carbon [11,17–21].

### Reclamation & Soil C and N

The effects of disturbance and reclamation on SOM pools have been most thoroughly studied in the context of mining [22–25]. Studies on oil and gas wellpads are less common, even though the scope of both disturbance and reclamation is different in oil and gas development compared to coal mining [19,26,27]. During establishment of wellpads, vegetation is stripped and the soil is scraped. Prior to 1983, soil scraping occurred only to level the wellpad site, and stockpiles were not re-spread after the wellpad was abandoned. After 1983, Onshore Oil and Gas Order Number 1 required stockpiling of topsoil for re-spreading during reclamation [28]. As a result, topsoil is now scraped and stored separately, then respread during reclamation. While these practices are intended to provide a suitable environment for seed germination and seedling growth, they also result in repeated soil disturbance.

The time needed for SOM to recover differs with the rate at which SOM associated with soil particles of different sizes is broken down; the labile pool, composed of SOM associated with coarse soil particles, is broken down relatively quickly and recovers on the scale of years to decades. The recalcitrant pool, composed of SOM associated with silt-clay particles, is broken down much more slowly and recovers on the scale of 1,000-10,000 years [6,8,29]. The constituents of SOM (organic carbon, inorganic nitrogen, and organic nitrogen) also recover at different rates, with available nitrogen generally recovering more quickly than carbon [8,30]. Furthermore, repeated soil disturbance, such as occurs during removal, stockpiling, and respreading, or such as that which occurs with cultivation, results in soil mixing, increased aeration, and, ultimately, increased SOM decomposition and macroaggregate disruption [6,8,11,19,31–33]. When combined, reduced plant inputs and increased soil disturbance result in SOM cycling out of the soil more quickly than it is incorporated, resulting in a net loss of carbon [1,33]. In comparison, soil that is maintained with minimal disturbance and maximum cover may show increased SOM concentration, slower turnover of macroaggregates and associated POM, and increased storage of organic carbon in recalcitrant microaggregates [5,10,11,20,34,35].

Much of the oil and gas development in the western United States takes place in big sagebrush (*Artemisia tridentata*) ecosystems, which now cover only ~56% of their original 60-million-hectare range [36–38]. Sagebrush ecosystems provide a variety of services and are especially important as habitat for both native and domesticated species, including Greater Sage Grouse, (*Centrocercus urophasianus*), pygmy rabbits (*Brachylagus idahoensis*), mule deer (*Odocoileus hemionus*), and pronghorn (*Antilocapra americana).* In addition, they are home to approximately 350 plant and animal species of conservation concern [38]. The ability of these systems to support healthy wildlife populations both during and after drilling is a concern among researchers and land managers, and it has received extensive study [36,39–42]. However, though good soil function is vital to successful recovery, the ability of reclamation to restore soil function has not received similar attention. Wyoming has one of the largest extents of sagebrush systems in the western United States, making oil and gas disturbance in this state a particular concern [43,44]. Prior to the 2015 downturn in drilling, it was estimated that up to 20,000 additional wells could be drilled in Wyoming by 2018 [45], resulting in extensive impacts to big sagebrush ecosystems in addition to those already occurring.

In this study we investigate how indicators of soil organic matter recovery differ over a chronosequence of sites disturbed between 7 and 91 years ago and seek to relate differences between these indicators for pairs of wellpads and undisturbed sites to reclamation activities. The goal of our study is to better understand how reclamation associated with oil and gas development affects carbon and nitrogen pools, as both carbon and nitrogen contribute heavily to soil organic matter. We have designed our study to answer 2 key questions:

1. How do wellpads differ from undisturbed areas with regard to carbon and nitrogen pools, and how do those differences change over time since wellpad abandonment?
2. How does reclamation influence differences between wellpads and undisturbed areas, and are these influences the same for C and N associated with coarse, fine, and silt-clay size soil particle classes?

We predicted that there would be little to no difference in coarse-associated C and N between wellpads and undisturbed plots, and that differences would increase as soil particle size, and therefore time to recovery after disturbance, decreased. We also predicted that the differences between wellpads and undisturbed plots would be smaller for unreclaimed wellpads than reclaimed wellpads due to reduced soil handling post-abandonment on unreclaimed wellpads. Overall, we expected to find the smallest differences in the coarse-associated C and N pools between unreclaimed wellpads and undisturbed sites, and the largest differences in the silt-clay-associated C and N pools between reclaimed wellpads and undisturbed sites.

## Methods

### Site Description and Selection

We selected sites south of Rock Springs, Wyoming, in the South Baxter Basin (lat:41.275484, lon:-109.363403). Soils in this area are Ustic Haplargids, moderately dry soils with poorly defined horizons [46]. The mean annual temperature in the area is 6°C and the mean annual precipitation is 22 cm.

We first identified approximately 30 wellpads in the area south of Rock Springs using the WOGCC website (http://wogcc.wyo.gov/) and limiting our search to only those wells which had been abandoned for at least 10 years. Once we located potential well sites, we worked with local land managers and employees of the WOGCC to identify and access the most suitable of the wells. Individual wellpad records often do not record details of reclamation, but after consultation with the WOGCC, Bureau of Land Management (BLM), Wyoming Department of Environmental Quality (WDEQ), researchers at the University of Wyoming, and environmental consultants, we designated wells plugged after 1983 as reclaimed (an effort was made to restore the site to its previous condition), and those before 1983 as unreclaimed (no attempt at restoration to initial conditions was made). We chose the cut-off date of 1983 because this was the year that reclamation was first mandated and enforced in Wyoming, and therefore the earliest we could expect reclamation to have occurred. In addition to the 1983 cut-off date, we sought physical evidence of reclamation occurring or not occurring. Because respreading topsoil was not required before reclamation was mandated, soil removed to level the sites was frequently left along one side of the wellpad when it was abandoned. These berms were still apparent on pre-1983 wellpads but non-existent on post-1983 wellpads. Furthermore, presence of planting lines from grass seedings were present on the wellpads abandoned in the mid-1980’s, corresponding to early reclamation regulations that did not limit the plant species used in reclamation. Sites abandoned as early as 1985, but no earlier, showed plentiful signs of being planted to crested wheatgrass, further supporting the use of 1983 as a reasonable cutoff year between reclaimed and unreclaimed wellpads. Ultimately, we selected 19 wellpad sites that were abandoned between 1943 and 2007, eight of which were reclaimed and eleven of which were not. Each site consisted of paired plots: a plugged and abandoned wellpad approximately 40 m in diameter, and an undisturbed control plot.

### Field Sampling

At each plot, we took sets of soil samples at three distances from the center of the plot using a hand-driven soil corer 7 cm in diameter. Each set of samples consisted of 2-3 sampling locations: understory (U, meaning beneath sagebrush), interspace (B, or between sagebrush), and on the mounds (M) formed by bunchgrasses, where applicable. Due to abundance of rocks in the local soil, we were generally limited to samples from 0-5 cm and 5-10 cm deep. However, where the soil was suitably clear of rocks, we also collected samples from 10-30 cm (Figure 1). Each sample was bagged separately and returned to the lab to be air-dried. In addition, we placed 3 pairs of cation-anion exchange resins, or “plant root simulator (PRS) probes” (Plant Root Simulator Probes, Western Ag Innovations, Saskatoon, SK, Canada) in each plot to estimate plant-available nitrate and ammonium. Probes remained buried in the top 5 cm of soil from October 2014 through May 2015, and were then returned to Western Ag Innovations for processing and analysis.

**Figure 1:**
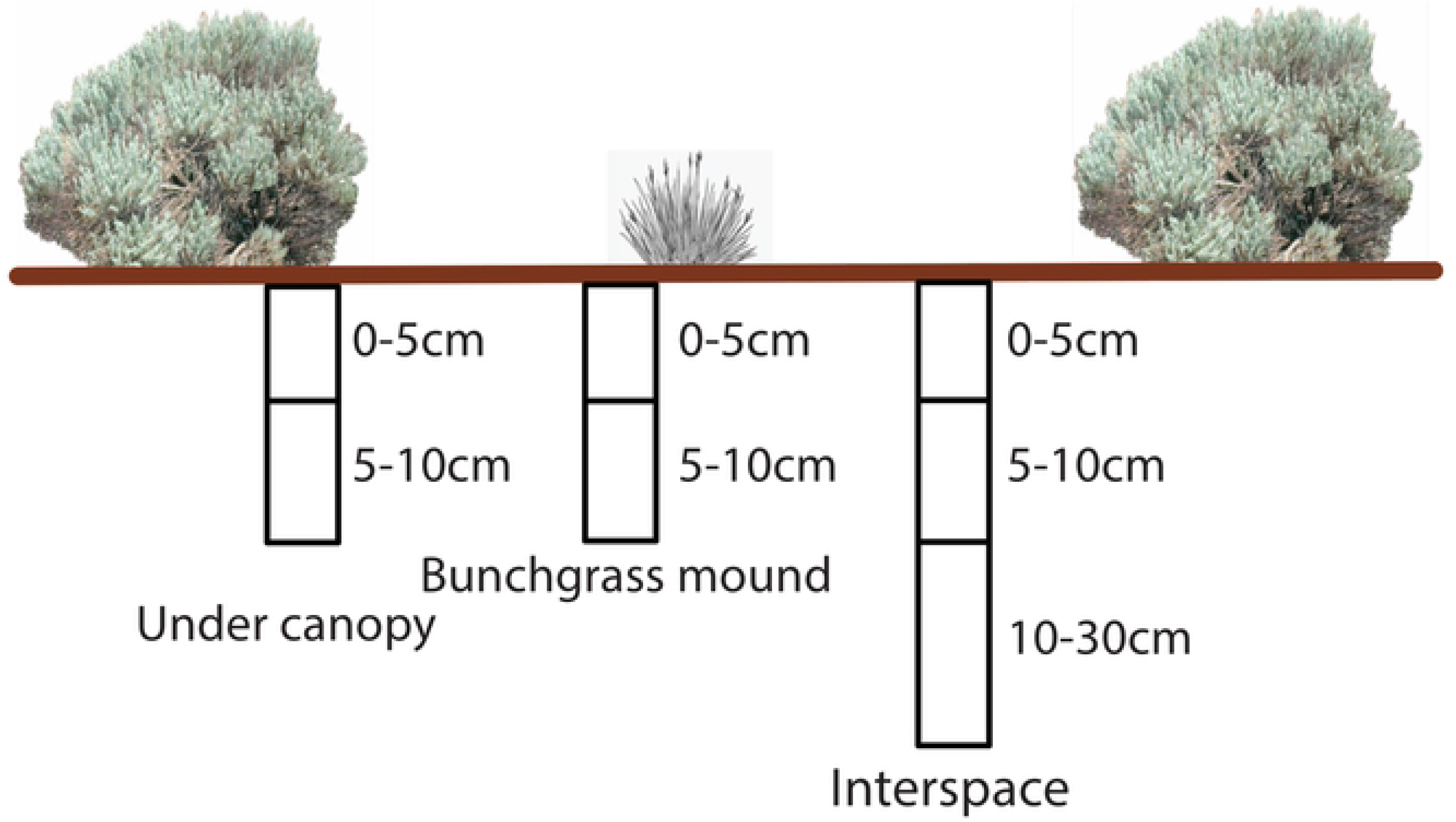
Sampling locations for each set of soil samples. 3 sets of soil samples were taken from each plot. In locations lacking shrubs, the “under canopy” sample was not taken.

### Soil Texture Analysis & Particle Size Separation

After we returned soil samples to the lab, we air-dried them and passed each through a 2 mm sieve to remove roots, leaves, and rocks. Each sieved sample was first analyzed for total C and N, and subsequently for C and N associated with three particle size classes: coarse (>250μm; least recalcitrant), fine (250-53 μm; intermediately recalcitrant), and silt-clay (<53 μm; most recalcitrant). We then composited 50 g of each sample by location and depth within each plot, resulting in 1 sample/depth/location/plot, or approximately 266 samples in total (7 per plot). We conducted texture analyses on the composited samples using Gee and Bauder’s hydrometer method [47].

To separate soils into these size classes, we used a size fractionation method [10,48]. Composited soil samples were submerged in de-ionized water on a 250 μm sieve and allowed to sit for 2 minutes. They were then sieved for 2 minutes, during which the screen of the sieve was raised and lowered 50 times, and any material left on the screen was transferred to a tin for drying. Material that passed through the sieve was poured onto a second sieve with a 53 μm screen and wet-sieved in the same manner as the first. We transferred the material on the sieve as well as the material that passed through the sieve into separate tins and placed the tins in drying ovens until all water had evaporated from the samples. We then homogenized the resulting dry soil samples and analyzed them for percent C and N on an elemental analyzer infrared mass spectrometer (EA-IRMS) at the University of Wyoming Stable Isotope Facility.

## Statistical Analyses

### Soil C and N at different depths and locations by treatment

We performed all of our statistical analyses in R [49] using multiple packages. For each plot, we received two sets of values for the PRS probes, corresponding to plant location (under canopy or interspace). To determine if there were differences between treatment, we used an ANOVA and, where appropriate, Tukey’s HSD post-hoc tests (‘lmerTest’ and ‘agricolae’ packages, respectively; [50,51]).

We separated our soil data into two groups for analysis of treatment differences among depths and locations. The first group (“surface soils”) contained all plant locations (interspace, under canopy, and bunchgrass mound) from 0-5 cm and 5-10 cm. The second group (“all depths”) contained all depths (0-5 cm, 5-10 cm, and 10-30 cm) at the interspace sampling location. To analyze for effect of treatment, each sample was assigned a treatment identification: undisturbed plot, unreclaimed wellpad, or reclaimed wellpad. We then used ANOVAs as well as Tukey’s HSD post-hoc tests to test for differences in C and N associated with either plant location (surface soils) or depth (all depths).

In addition to ANOVAs, we used hierarchical partitioning to identify the relative importance of each of a series of variables (plant location, depth, years since abandonment, and reclamation treatment) to variation in C and N (‘relaimpo’ package; [52]). Hierarchical petitioning calculates the contribution of each predictor variable to the variance of the dependent variable by evaluating its effect in every possible model and reporting the mean of these effects as the percent contribution to variance. In doing so, it helps alleviate the problem of multicollinearity that is found in some other modeling approaches used to identify relative importance of predictor variables [53,54].

### Differences over time for C and N on wellpads and undisturbed plots

To test for effects of time and reclamation on C and N recovery, we subtracted the mass of C or N on the wellpad from the mass of C or N on the corresponding undisturbed plot. We then performed regression analyses to identify any significant relationships between time (years since wellpad abandonment) and changes in the difference between wellpads and undisturbed areas. We defined a decrease in the difference between the wellpads and undisturbed areas as a trajectory towards recovery. We also performed regressions separately for each wellpad treatment (reclaimed or unreclaimed) and used the slope of the regression to identify whether the rate of recovery was greater on reclaimed than unreclaimed wellpads. We used the same procedure to test for the effects of time and treatment on recovery of total NH_4_ + NO_3_.

## Results

### C and N on wellpads vs. undisturbed plots

Plant location (under a shrub canopy, in the bare interspace between plants, or directly under a bunchgrass mound) and treatment (undisturbed plot, unreclaimed wellpad, or reclaimed wellpad), as well as their interactions, had the largest effect on soil carbon and nitrogen (Table 1). The mass of C or N was generally greater under canopies or on bunchgrass mounds than in the interspace, and generally decreased with sample depth (Figure 2). Both layers of soil from bunchgrass mounds contained significantly more N on undisturbed plots than they did on wellpads, while the topmost layer of soil from interspaces had significantly less C on undisturbed plots than it did on wellpads. Plant-available N as measured using the PRS probes did not differ between sites or locations.

**Table 1:**
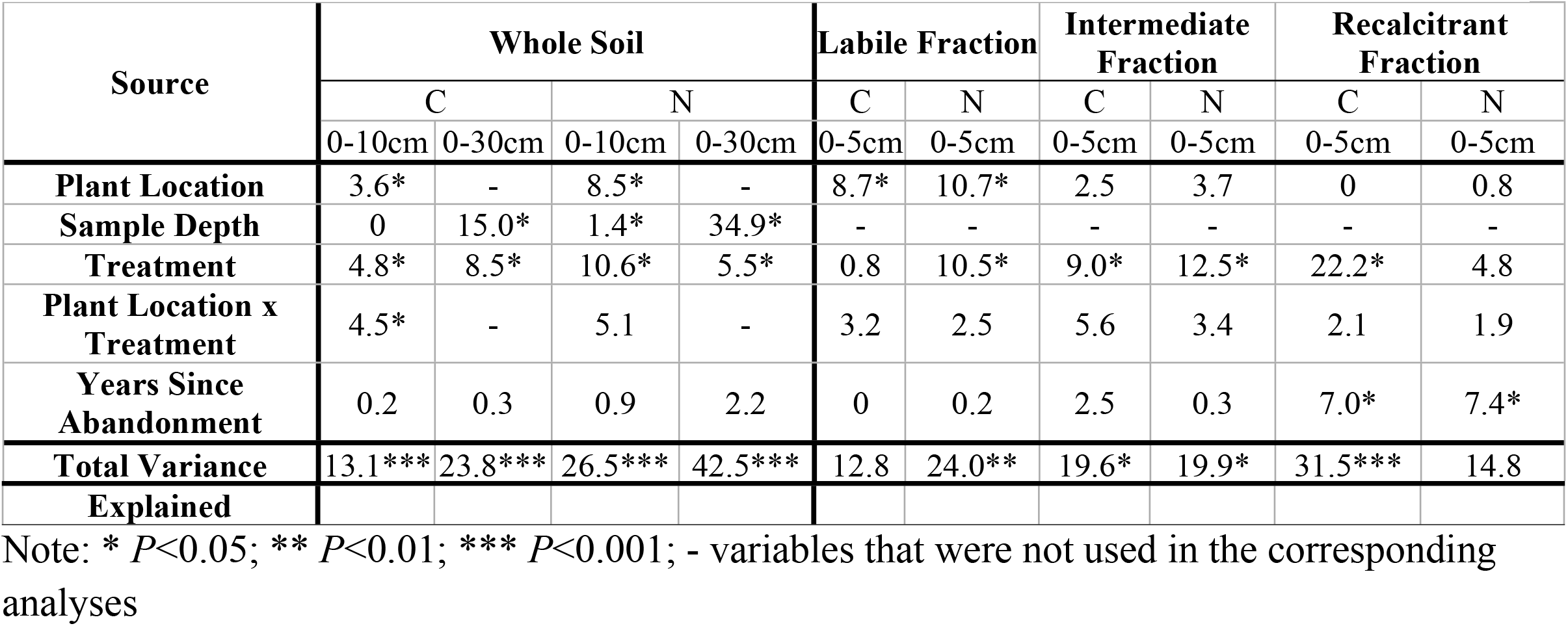
Percent of total variance in mass of carbon and nitrogen explained by predictor variables in ANOVAs of data from 19 wellpad sites

**Figure 2:**
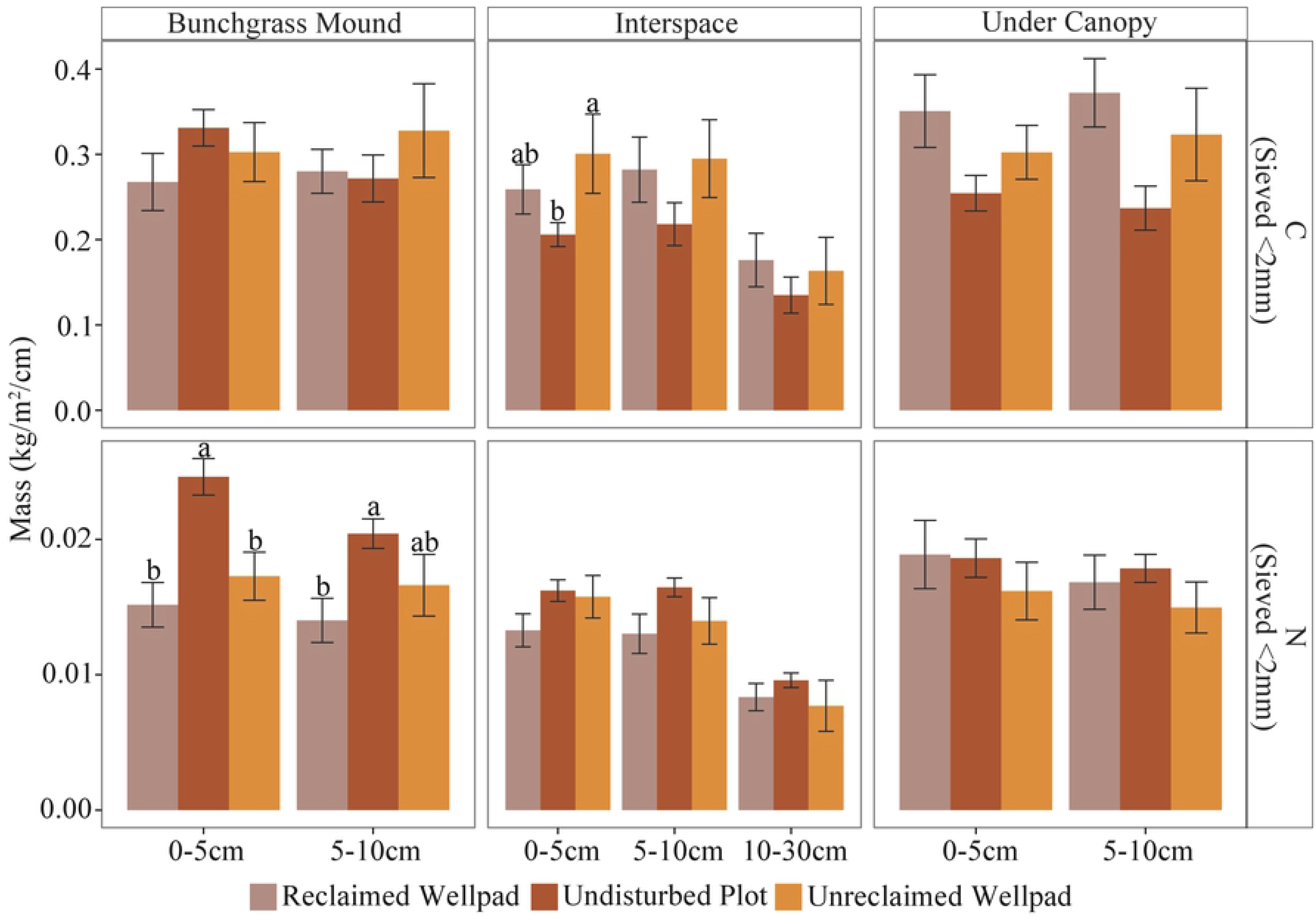
Mass of C and N at three depths and three locations on reclaimed or unreclaimed wellpads and undisturbed plots. Significant differences between groups are indicated by letters. Errors bars represent 1 standard error above and below the mean.

Interspace soil contained less C and N than plant canopy soils, regardless of the associated soil particle size and whether the site was a wellpad or undisturbed plot. Likewise, regardless of whether the site was a wellpad or undisturbed, coarse particles were usually associated with the highest mass of both N and C, while fine particles were usually associated with the lowest (Figure 3).

**Figure 3:**
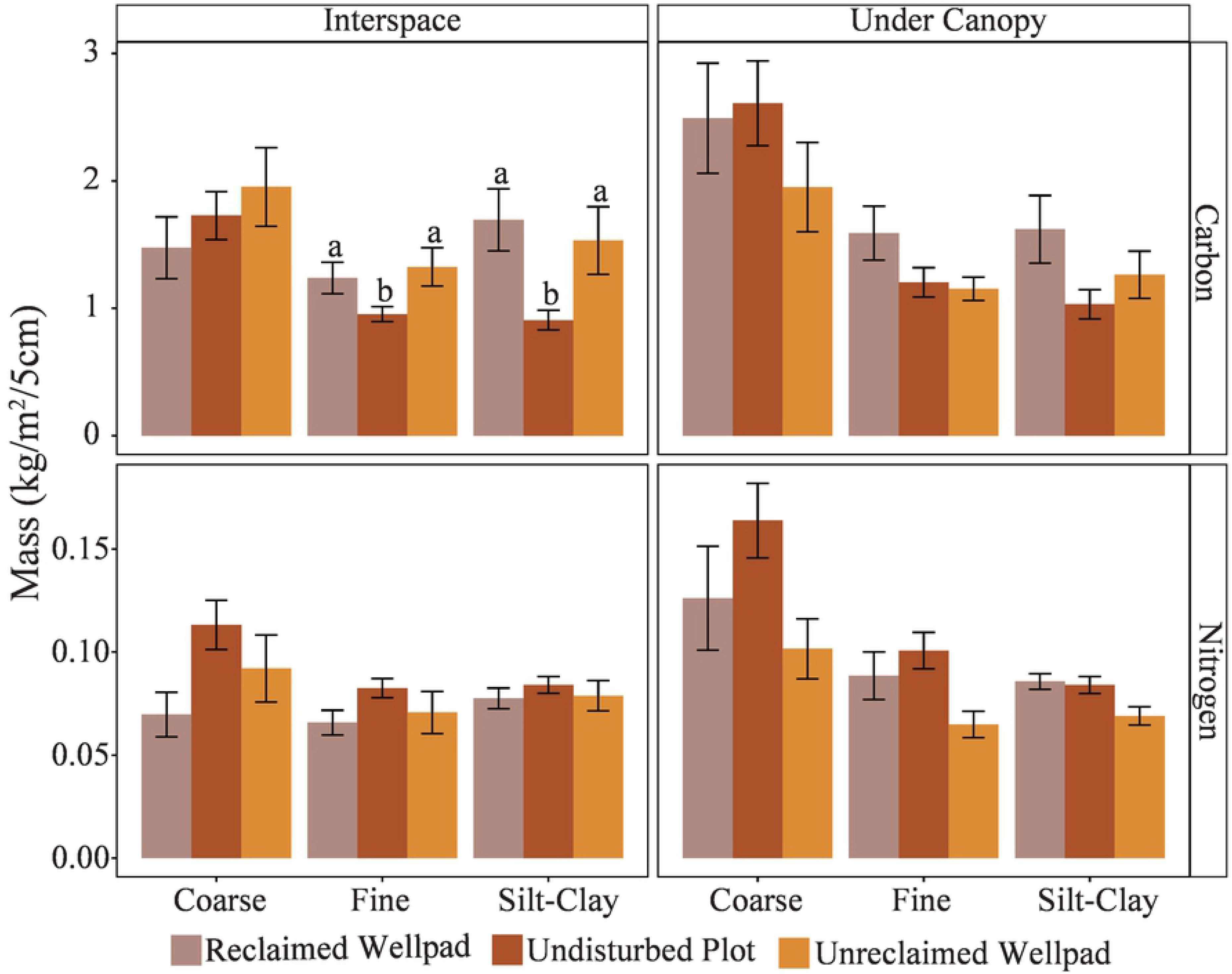
Mass of C and N associated with three soil particle size fractions at two sampling locations on reclaimed or unreclaimed wellpads and undisturbed plots. Significant differences between groups are indicated by letters. Errors bars represent 1 standard error above and below the mean.

There was significantly less C associated with both fine and silt-clay particles from undisturbed plots than from wellpads. As particle size decreased, treatment became more important in explaining mass of C, and plant location became less important in explaining mass of both C and N (Table 1). Nitrogen associated with silt-clay particles differed less with position than did nitrogen associated with coarse or fine particles (Figure 3).

Time since abandonment did not have a significant effect on differences in mass of C between wellpads and undisturbed plots, regardless of depth or plant location. Differences in nitrogen, however, decreased significantly in surface soils at all plant locations as time since abandonment increased, indicating partial recovery (Figure 4). Differences between wellpads and undisturbed plots also decreased significantly for interspace C and N associated with coarse particles, and for N associated with silt-and-clay particles, as time since abandonment increased (Figure 5).

**Figure 4:**
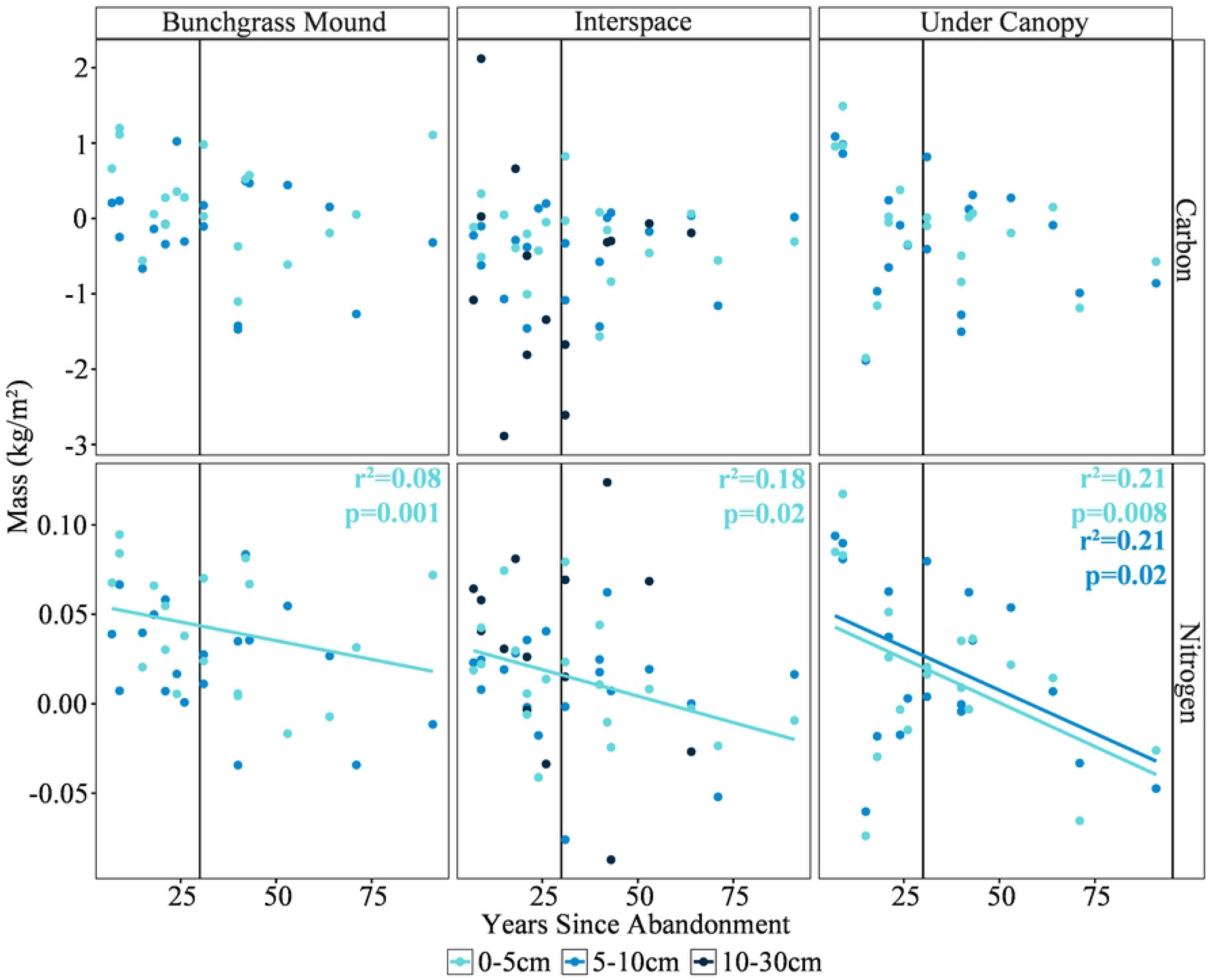
Difference in mass of C and N between wellpads and undisturbed sites vs. time since abandonment at three depths and three sampling locations. Positive numbers indicate that mass on the wellpad was lower than mass on the undisturbed plot, and negative numbers indicate the opposite relationship. P-values and r^2^ values are noted where the correlation between time and difference was significant.

**Figure 5:**
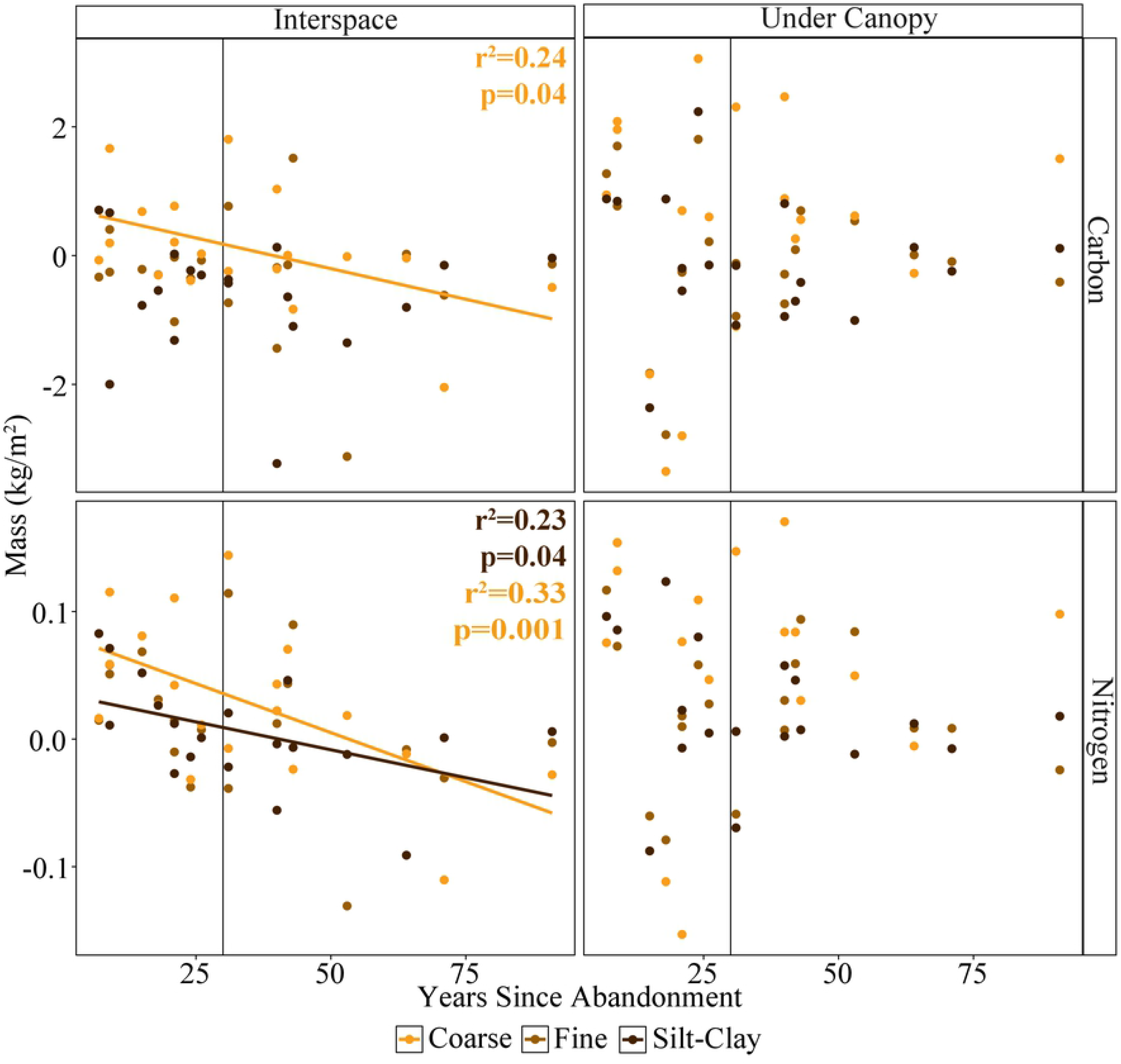
Difference in mass of C and N vs. time since wellpad abandonment in three soil particle size fractions at two sampling locations. Positive numbers indicate that mass on the wellpad was lower than mass on the undisturbed plot, and negative numbers indicate the opposite relationship. P-values and r^2^ values are noted where the correlation between time and difference was significant.

### Effects of wellpad reclamation on C and N

Regardless of plant location, unreclaimed and reclaimed wellpads did not differ significantly from each other in mass of C or N (Figure 2). Likewise, mass of C and N did not differ significantly between reclaimed and unreclaimed wellpads for any particle size class.

Wellpads became more like undisturbed plots over time for some individual depths and plant locations, but our regressions found no consistent relationship between reclamation and C or N recovery at any depth or plant location. There was likewise no relationship between reclamation and recovery on the combined depth samples, though the difference in mass of N between reclaimed wellpads and undisturbed areas did decrease for interspace samples (Figure 6). Differences in interspace N between unreclaimed wellpads and undisturbed areas decreased for all particle sizes as time since abandonment increased, but we did not note any other relationships between reclamation and mass of C or N associated with the other particle sizes and plant locations.

**Figure 6:**
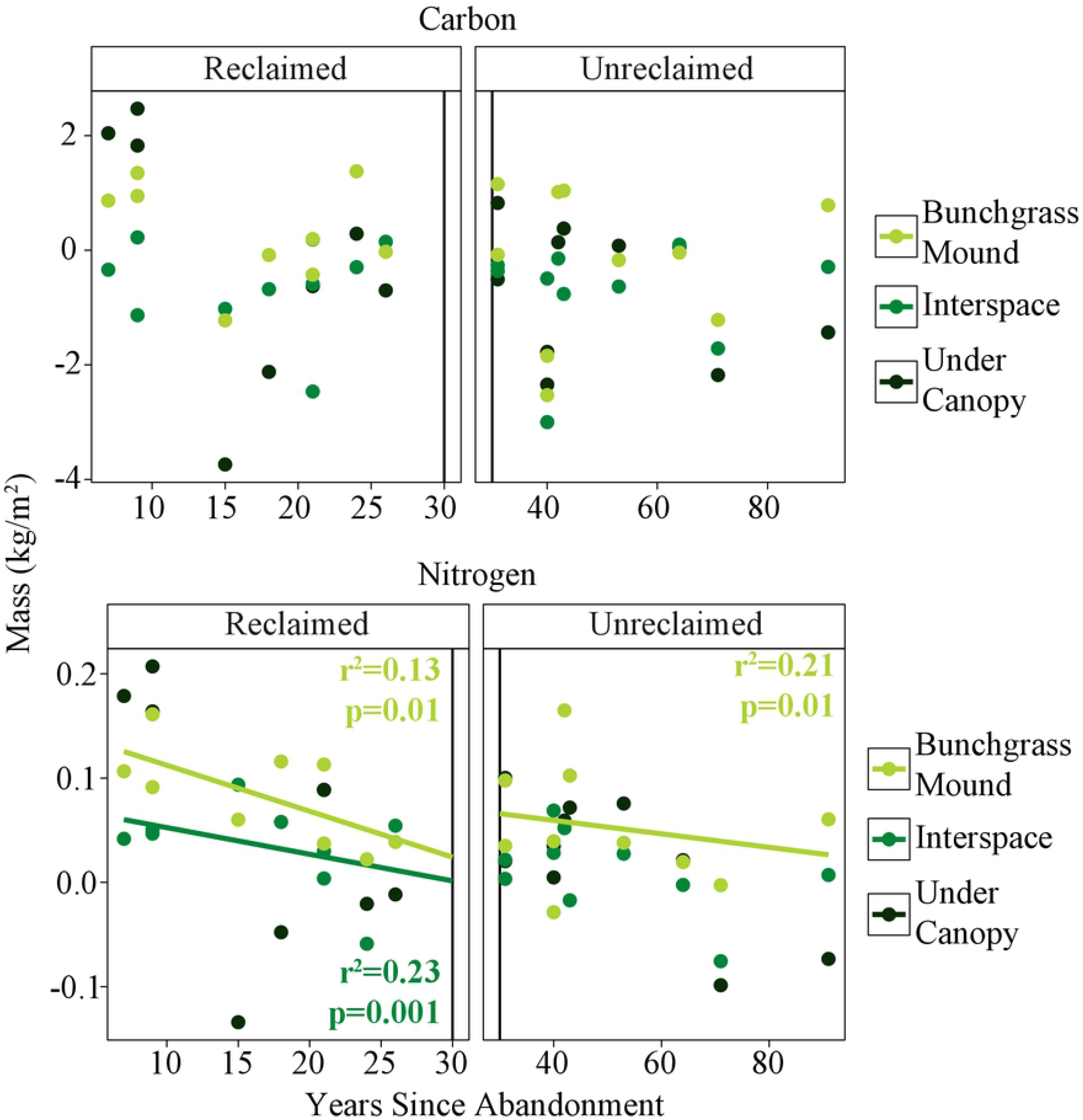
Difference in whole soil core mass of N and C vs. time since wellpad abandonment at three sampling locations. Positive numbers indicate that mass on the wellpad was lower than mass on the undisturbed plot, and negative numbers indicate the opposite relationship. P-values and r^2^ values are noted where the correlation between time and difference was significant.

## Discussion

We found that the most important factors in determining soil C and N recovery were not reclamation treatment but plant location, soil depth, and time since abandonment. Overall, there was little difference in C and N between reclaimed and unreclaimed wellpads, which indicates that reclamation did not have a positive or negative effect on the recovery of these carbon and nitrogen pools or the SOM to which they contribute. Our prediction that differences between treatments would be more pronounced in the silt-clay particle size than the coarse particle size was not supported by our findings. Rather, the opposite occurred, with the largest amount of variation in the coarse particle size and the smallest in the silt-clay particle size. In addition, because there was very little difference between unreclaimed and reclaimed wellpads, we cannot conclude whether treatment effects changed with increasingly fine particle sizes.

For both C and N, plant location decreased in importance with fineness of soil particle size. Both C and N associated with coarse particle sizes were most sensitive to plant location, while C and N associated with silt-and-clay sizes were least sensitive. Previous work has shown that coarse fractions of SOM are very sensitive to inputs and outputs [8], including their quality (C:N ratio) and quantity [55]. The importance of plant location to these fractions may reflect this increased sensitivity. Conversely, very fine silt-and-clay fractions are less sensitive to inputs and outputs than coarse fractions [56], and less likely to reflect small-scale heterogeneity of inputs due to plants. This should be especially true in surface soils, which are more vulnerable to mixing between interspace and canopy positions.

Carbon was generally higher on wellpads than undisturbed adjacent areas, which is contrary to most literature concerning disturbance and soil organic matter. In most other studies, disturbance resulted in a loss of soil organic matter, including carbon [6,8,11,19,31–33]. There may, however, be several factors causing higher carbon amounts on wellpads than undisturbed plots in our study. First, on at least some of the sites, soil was stockpiled, seeded as part of interim reclamation, and re-spread during final reclamation. The plants seeded on stockpiles are usually quick-growing grasses with extensive root systems that stabilize the soil against erosion. These grasses are not removed when the soil is respread, and are therefore incorporated back into the soil. Furthermore, grasses form a large proportion of reclamation seed mixes. Grass roots in other systems have been shown to contribute significantly to spatial variability in total SOM due to their large biomass [56]. In dryland ecosystems, previous research has shown that it is plant presence, and not plant attributes, that contribute most significantly to soil nutrients in areas with patchy cover [55]. Extensive rooting systems as well as increased litter input and the capability of bunchgrasses to form small ‘nutrient islands’ [57–59] have been suggested previously to explain a relatively greater increase in C than N on sites in sagebrush steppe where shrubs had been removed [60]. This combination of factors (plant presence, large root biomass, increased litter input, and formation of nutrient islands) is a possible explanation for the disproportionate increase in C over N on our wellpad sites.

We used a chronosequence approach to address questions about soil organic matter recovery on disturbed sites in a big sagebrush ecosystem. Chronosequences have inherent limitations in that they are space-for-time substitutions, and therefore do not measure one site over time, but many sites from across a long time period [61,62]. They are therefore not appropriate for answering questions about systems that are frequently disturbed or follow unpredictable trajectories [61]. In the case of our study, we assumed that all sites were on similar trajectories and that soil organic matter recovers in a predictable manner. Our inability to say with 100% certainty that this is the case is a limitation of our study, and of all chronosequence studies [61], but does not preclude addressing general questions. An additional limitation in our study was the conflation of time since abandonment and an individual site’s treatment during oil and gas development, or reclamation status. However, if reclamation was having a significant effect on soil organic matter pools, we would expect these effects to be identifiable in our treatments, and our findings are consistent with previous research [26]. Furthermore, while this conflation may have affected comparisons between reclaimed and unreclaimed wellpads, it did not affect our ability to answer questions about how wellpad disturbance in general has affected C and N relative to undisturbed areas.

We found that disturbance resulted in lower N on wellpads than on undisturbed sites. This is counter to the results of Avirmed et al. (2015), who studied old, unreclaimed wellpads that had not experienced respreading of stockpiled topsoil, and found no difference in soil organic matter between those wellpads and adjacent undisturbed areas. More importantly, we found that reclaimed sites were not different from unreclaimed sites, and most importantly, that the rate at which they recovered over the time since they were abandoned was not significantly faster relative to sites where reclamation was not attempted. This suggests that reclamation did not cause a significant gain over natural processes where rate of recovery was concerned, with important implications for oil and gas development across the sagebrush region. Successful reclamation of a single wellpad has been estimated to cost $16,000-$17,000. If the first effort is not successful, subsequent attempts can cost an additional $20,000 to $40,000. The capital investment represented by reclamation procedures necessitates that they minimize cost while maximizing effectiveness. It is therefore important to better clarify which aspects of reclamation lead to improved soil recovery, and which do not.

